# HIGH FIDELITY DETECTION OF CROP BIOMASS QTL FROM LOW-COST IMAGING IN THE FIELD

**DOI:** 10.1101/150144

**Authors:** Darshi Banan, Rachel Paul, Max Feldman, Mark Holmes, Hannah Schlake, Ivan Baxter, Andrew D.B. Leakey

**Affiliations:** University of Illinois Urbana Champaign 1402 IGB 1206 W Gregory Dr Urbana, IL 61801 USA; Donald Danforth Plant Science Center 975 North Warson Road St. Louis, MO 63132 USA; USDA-ARS, Donald Danforth Plant Science Center 975 North Warson Road St. Louis, MO 63132 USA

## Abstract

Above-ground biomass production is a key target for studies of crop abiotic stress tolerance, disease resistance and yield improvement. However, biomass is slow and laborious to evaluate in the field using traditional destructive methods. High-throughput phenotyping (HTP) is widely promoted as a potential solution that can rapidly and non-destructively assess plant traits by exploiting advances in sensor and computing technology. A key potential application of HTP is for quantitative genetics studies that identify loci where allelic variation is associated with variation in crop production. And, the value of performing such studies in the field, where environmental conditions match that of production farming, is recognized. To date, HTP of biomass productivity in field trials has largely focused on expensive and complex methods, which – even if successful – will limit their use to a subset of wealthy research institutions and companies with extensive research infrastructure and highly-trained personnel. Even with investment in ground vehicles, aerial vehicles and gantry systems ranging from thousands to millions of dollars, there are very few examples where Quantitative trait loci (QTLs) detected by HTP of biomass production in a field-grown crop are shown to match QTLs detected by direct measures of biomass traits by destructive harvest techniques. Until such proof of concept for HTP proxies is generated it is unlikely to replace existing technology and be widely adopted. Therefore, there is a need for methods that can be used to assess crop performance by small teams with limited training and at field sites that are remote or have limited infrastructure. Here we use an inexpensive and simple, miniaturized system of hemispherical imaging and light attenuation modeling to identify the same set of key QTLs for biomass production as traditional destructive harvest methods applied to a field-grown Setaria mapping population. This provides a case study of a HTP technology that can deliver results for QTL mapping without high costs or complexity.

## INTRODUCTORY PARAGRAPH

Above-ground biomass production is a key target for studies of crop abiotic stress tolerance, disease resistance and yield improvement. However, biomass is slow and laborious to evaluate in the field using traditional destructive methods^1^. High-throughput phenotyping (HTP) is widely promoted as a potential solution that can rapidly and non-destructively assess plant traits by exploiting advances in sensor and computing technology^2^. A key potential application of HTP is for quantitative genetics studies that identify loci where allelic variation is associated with variation in crop production. And, the value of performing such studies in the field, where environmental conditions match that of production farming, is recognized^3^. To date, HTP of biomass productivity in field trials has largely focused on expensive and complex methods, which – even if successful – will limit their use to a subset of wealthy research institutions and companies with extensive research infrastructure and highly-trained personnel. Even with investment in ground vehicles, aerial vehicles and gantry systems ranging from thousands to millions of dollars, there are very few examples where Quantitative trait loci (QTLs) detected by HTP of biomass production in a field-grown crop are shown to match QTLs detected by direct measures of biomass traits by destructive harvest techniques^4^. Until such proof of concept for HTP proxies is generated it is unlikely to replace existing technology and be widely adopted. Therefore, there is a need for methods that can be used to assess crop performance by small teams with limited training and at field sites that are remote or have limited infrastructure. Here we use an inexpensive and simple, miniaturized system of hemispherical imaging and light attenuation modeling to identify the same set of key QTLs for biomass production as traditional destructive harvest methods applied to a field-grown *Setaria* mapping population. This provides a case study of a HTP technology that can deliver results for QTL mapping without high costs or complexity.

## TEXT

Current rates of yield gain are unlikely to meet the projected demands of global population growth and development^5,6^. High-throughput phenotyping (HTP) techniques rapidly evaluate plant performance and leverage advances in genotyping^7–9^, the development of mapping populations^10–12^, and the design and analysis of quantitative genetic experiments to ultimately develop a predictive understanding of genotype-to-phenotype relationships^13,14^. This understanding enables an accelerated and more targeted approach to crop improvement^2,3^. Biomass production generates both the calories that are partitioned towards food consumption and the raw feedstock used for carbon efficient biofuels^15^. Biomass productivity per unit ground area must be maximized to ensure profitability and avoid displacement or disturbance of natural ecosystems^16^. However, biomass is a complex trait that is difficult to assess in the field and usually is measured by destructive harvest^1^. We addressed this challenge by testing the ability of hemispherical imaging to identify genomic regions associated with above-ground biomass production. The successful application of hemispherical imaging was evaluated with respect to its strong phenotypic correlation and high degree of QTL co-localization with directly validated destructive harvest traits.

Numerous remote sensing methods have been demonstrated to correlate with destructive measures of biomass production and can be considered proxy measurements^17–20^. These include multispectral and hyperspectral indices of radiation reflected from crop canopies, as well as measures of canopy light distribution. Often, a combination of sensor outputs and additional processing techniques such as regression, inverse modeling, and multivariate analysis are required to produce relevant phenotypes^1,21^. Tanger et al (2017) used a combination of tractor-mounted multispectral reflectance and ultrasonic sensors to detect manually validated QTL associated with biomass in rice^4^. Similar measurements have been deployed in other field-grown crops using ground vehicles, aerial vehicles and gantries requiring investment in equipment that often exceeds $100,000s - $1,000,000s^22,23^. In addition to the expense of HTP equipment, many techniques under development require extensive research infrastructure, permits (e.g. flight authorization) and complex data analyses. Highly trained personnel are consequently needed to support both data acquisition and analysis. Unfortunately, these factors combine to mean that the majority of HTP techniques can only feasibly be used by large research intuitions and companies. And, even in those organizations deployment of HTP has to be limited to a few high priority projects. Cheap methods of HTP that rely on simpler technology could greatly increase how widely HTP is adopted, and support work in a broader diversity of environmental conditions and crops outside of the major growing regions of the world’s staple crops.

Hemispherical imaging captures the geometry of sky openings and models the attenuation of solar radiation by the canopy to estimate canopy properties, such as Leaf Area Index (LAI; leaf area per unit ground area)^24,25^. The ability to account for the influence of both stems and leaves allows its use for estimating Plant Area Index (PAI; above-ground plant tissue area per unit ground area) in herbaceous systems^26^. Canopy properties estimated from hemispherical images use mechanistic and biophysical models rather than reliance on statistical relationships between sensor and subject. Therefore, they should be less context dependent and more widely applicable to different crops, growing conditions, and management practices than other methods that require a training model to relate remotely sensed data to traditional measures of crop productivity^27^ (e.g. Busemeyer et al 2013). Traditionally, hemispherical photography equipment is tall, bulky, and not suited to crop HTP. In this study a miniature remotely triggered digital camera designed for point-of-view action sport videography (GoPro Hero 3+) was modified with a miniature hemispherical lens and mounted to a custom-built self-leveling gimbal (Fig. 1 a-c). The resulting system was small enough to fit between tight crop rows and below the crop canopy.

**Figure 1.**
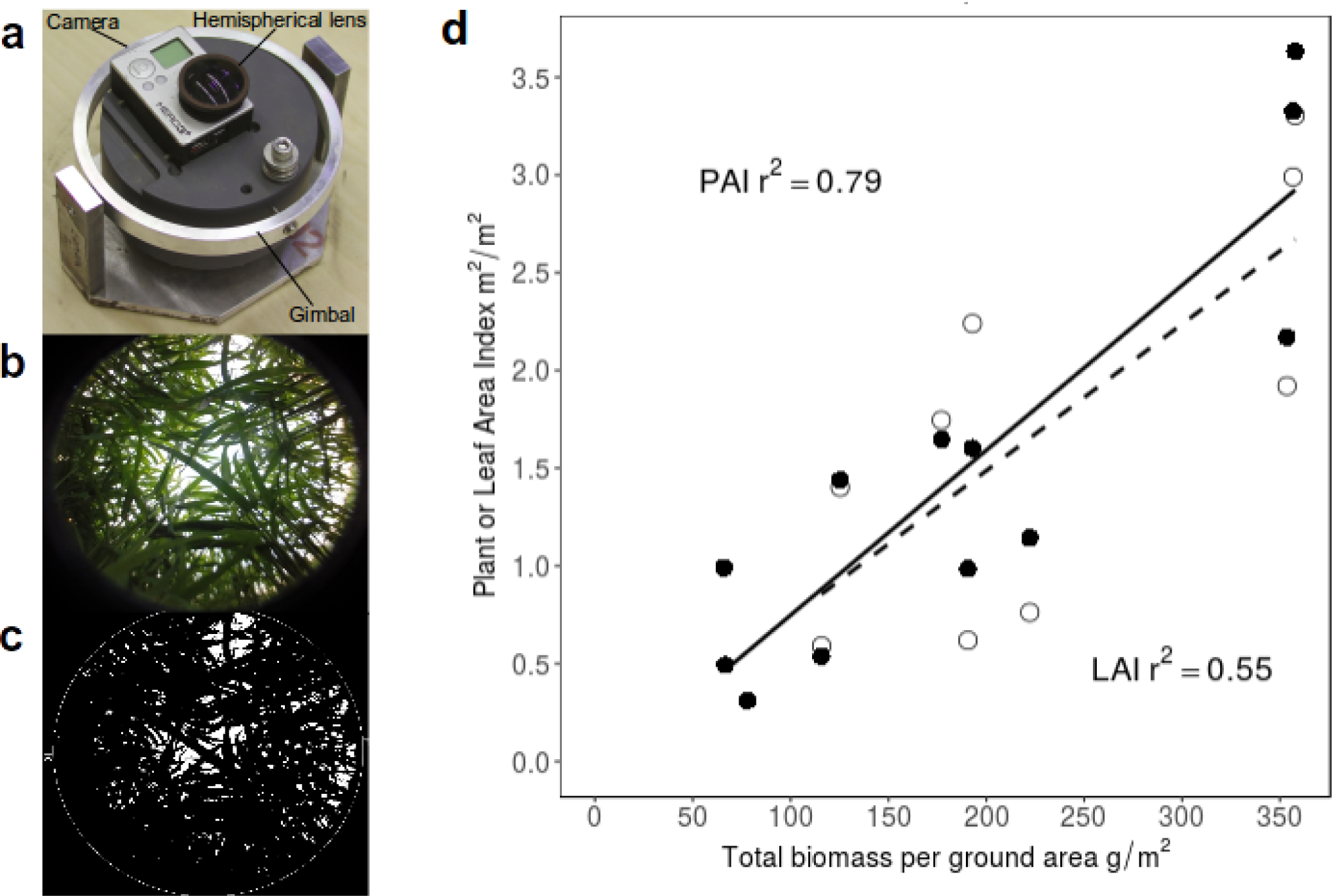
Customized hemispherical imaging system application to high throughput phenotyping of above-ground biomass production. **a,** A hemispherical lens fitted on a GoPro Hero3+ digital camera and mounted on a self-leveling gimbal. The camera unit had maximum dimensions of 5.8*4*3.7 cm and the full system was 11*15*13.3 cm. **b,** The system was used to capture fully hemispherical images of a plant canopy. **c,** Images were thresholded for analysis and estimation of Plant Area Index (PAI) using Delta-T Hemiview software. **d,** The resulting PAI estimates were correlated to total biomass (filled symbols, solid line) and compared to that between Leaf Area Index estimated from destructive harvest and total biomass (open symbols, dashed line). Measurements made on parent lines A.10, B.100 and phenotypically intermediate RIL#161 together represent a diversity of growth habit and morphology seen across the population. Symbols correspond to single plots from which all images and measurements were collected 38, 44, 52, and 60 days after sowing. Correlation r^2^ values are reported for both measurements.

An F7 recombinant inbred line (RIL) mapping population with 186 genotypes generated from a cross between the cultivated *Setaria italica* and its weedy ancestor *Setaria viridis* provides an ideal platform for assessing the ability of hemispherical imaging to detect QTL related to above-ground biomass production due to its wide diversity of morphologies and multifold variation in biomass production^7,11^. As a model C_4_ grass emerging as a tool for systems-level biology, the genus *Setaria* has the advantage of being closely related to C_4_ grass food and fuel crops such as maize, sorghum, miscanthus, and switchgrass, while having a smaller stature, faster life cycle, and diploid genome^28^.

First, a validation experiment determined whether the customized imaging system was suitable for HTP in *Setaria*. *Setaria viridis, Setaria italica,* and the phenotypically intermediate RIL #161 derived from crossing these two species were used as test material because they vary widely in canopy architecture and rate of biomass production. Each genotype was grown in eight replicated plots of which four were randomly chosen for measurement. An independent plot for each genotype was selected for collection of both hemispherical images and destructive canopy and biomass harvest measures on four dates distributed across the growing season to generate a wide range of canopy closures and biomasses with which to evaluate hemispherical imaging. Notably, total above-ground biomass correlated more strongly with PAI (r^2^=0.79) than it did with destructively measured LAI (r^2^=0.55, Fig. 1d). This highlights the ability of hemispherical imaging to robustly assess the total amount of plant tissue over an area of ground across a diversity of short, herbaceous grass canopies in a non-destructive manner. Data is easy to acquire and analyze since the camera, lens and analysis software are all commercially available.

Next, the ability of hemispherical imaging to detect QTLs for biomass production was tested in a *Setaria* F7 RIL mapping population. 186 RILs were planted in a randomized design with six check plots for each parent. Hemispherical images, manually measured morphological traits, and destructive harvest weight data were collected from the same plots of each genotype.

Results from manual measures of developmental, architecture, and biomass production traits showed that the segregating population was phenotypically diverse for a comprehensive set of destructive harvest traits assessed at maturity, including total above-ground biomass, tiller number, and height (Table S1).

Principal component analysis (PCA) was performed on directly measured traits – leaf mass, panicle mass, stem mass, branch number, clump spread, culm height, tiller height, days-after-sowing until panicle emergence, and reproductive-to-vegetative mass ratio – to simplify the description of plant performance relative to hemispherical imaging estimates (Fig. 2). The first three principal components (PCs) with eigenvalues greater than 1.00 together explain 76% of the variation in the dataset. The trait loadings based on eigenvectors in each of these three orthogonal PCs appear to describe three plant growth components: (1) biomass production, (2) bushiness, and (3) partitioning of biomass to vegetative versus reproductive structures. Days-after-sowing until panicle emergence varied significantly within the RIL population, loading moderately in both PC1 and PC3, but not strongly in any single PC. PC1 accounted for 44% of overall variation and approximates above-ground biomass production based on strong loadings for culm height, tiller height, leaf mass, panicle mass, and stem mass. PC2 accounted for 20% of variation and approximates plant bushiness based on strong loadings for clump spread, branch number, and tiller number. PC3 accounted for 12% of variation and approximates the partitioning of biomass to vegetative versus reproductive structures based on strong loadings for panicle mass and reproductive-to-vegetative mass ratio.

**Figure 2.**
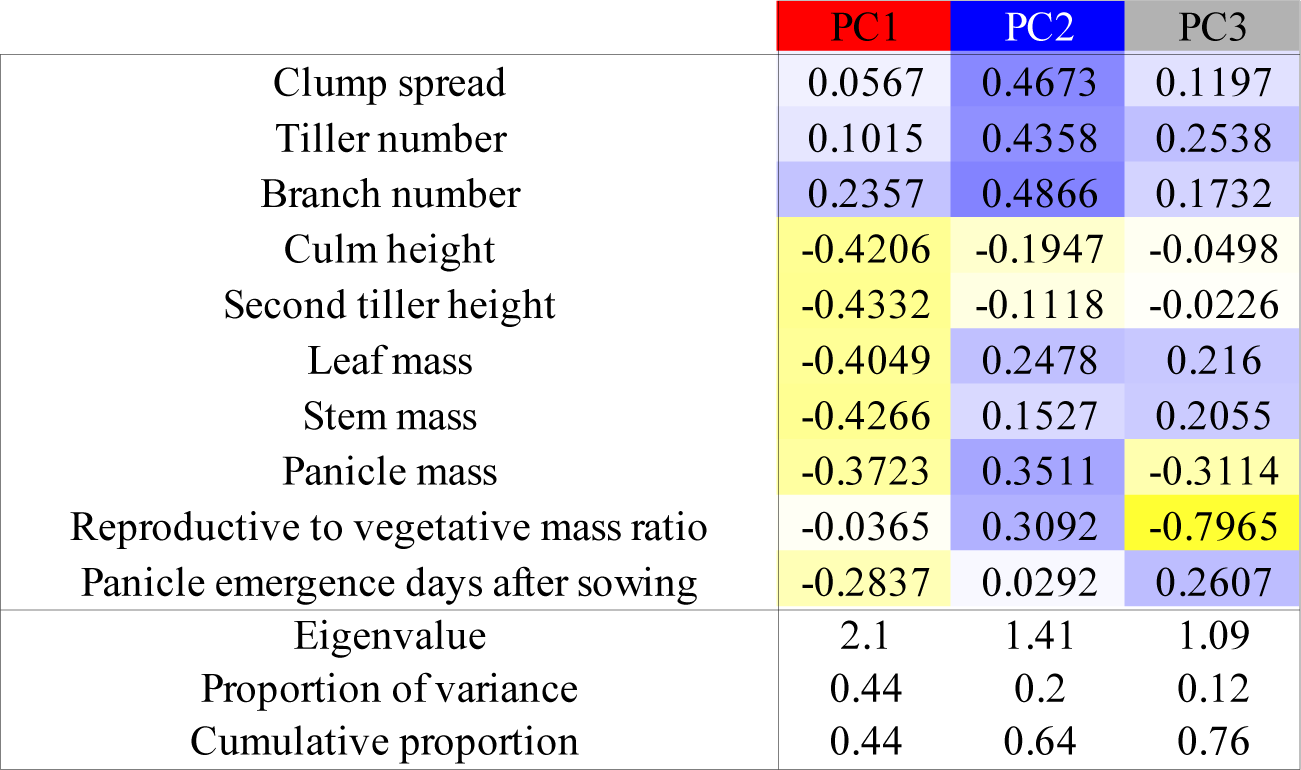
Principal component analysis of directly measured biomass traits. Trait loadings based on eigenvectors (color-coded by direction and magnitude, blue: positive, yellow: negative) in each principal component (PC) appear to describe three orthogonal processes: biomass production (PC1), bushiness (PC2), and partitioning of biomass to vegetative versus reproductive structures (PC3). The individual and cumulative contributions of each PC are reported.

These apparent descriptions, while not definitive, are biologically intuitive and provide a framework for interpreting the genetic and phenotypic attributes of a plant in the field. Correlation analysis of all traits measured shows that PAI correlated positively and strongly with total mass, vegetative mass, and with the other traits that loaded strongly into PC1 to describe biomass production (Fig. 3).

**Figure 3.**
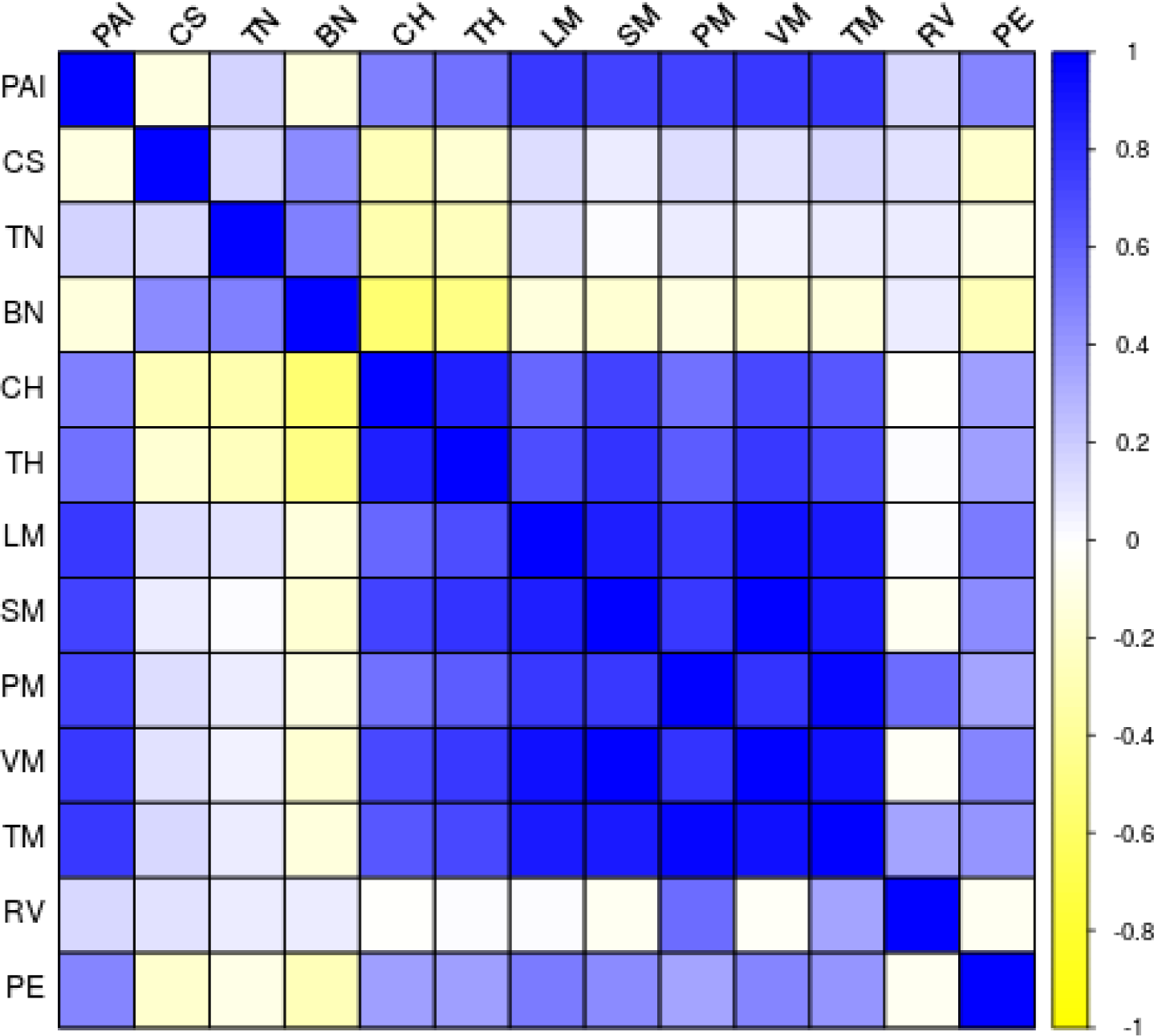
Correlation matrix of PAI and directly measured biomass traits. PAI correlates strongly and positively with traits associated with biomass production. Blue, yellow, and white cells represent positive, negative, and no correlation, respectively, between traits based on Pearson correlation coefficient values. PAI=plant area index, CS=clump spread, TN=tiller number, BN=branch number, CH=culm height, TH=second tiller height, LM=leaf mass per m^2^ ground, SM=stem mass per m^2^ ground, PM=panicle mass per m^2^ ground, VM=vegetative mass per m^2^ ground, TM=total mass per m^2^ ground, RV=reproductive to vegetative mass ratio, PE=panicle emergence days after sowing.

Quantitative trait loci analysis detected 53 significant QTL across 12 traits (Table S2). The identified loci clustered in groups corresponding with the segregation of traits into the three different PCs (Fig. 4). This suggests that the variation underlying the separation of PCs is driven by genetic programs related to biomass production, bushiness, and partitioning of biomass to vegetative versus reproductive structures rather than uncontrolled physiological or environmental factors.

**Figure 4.**
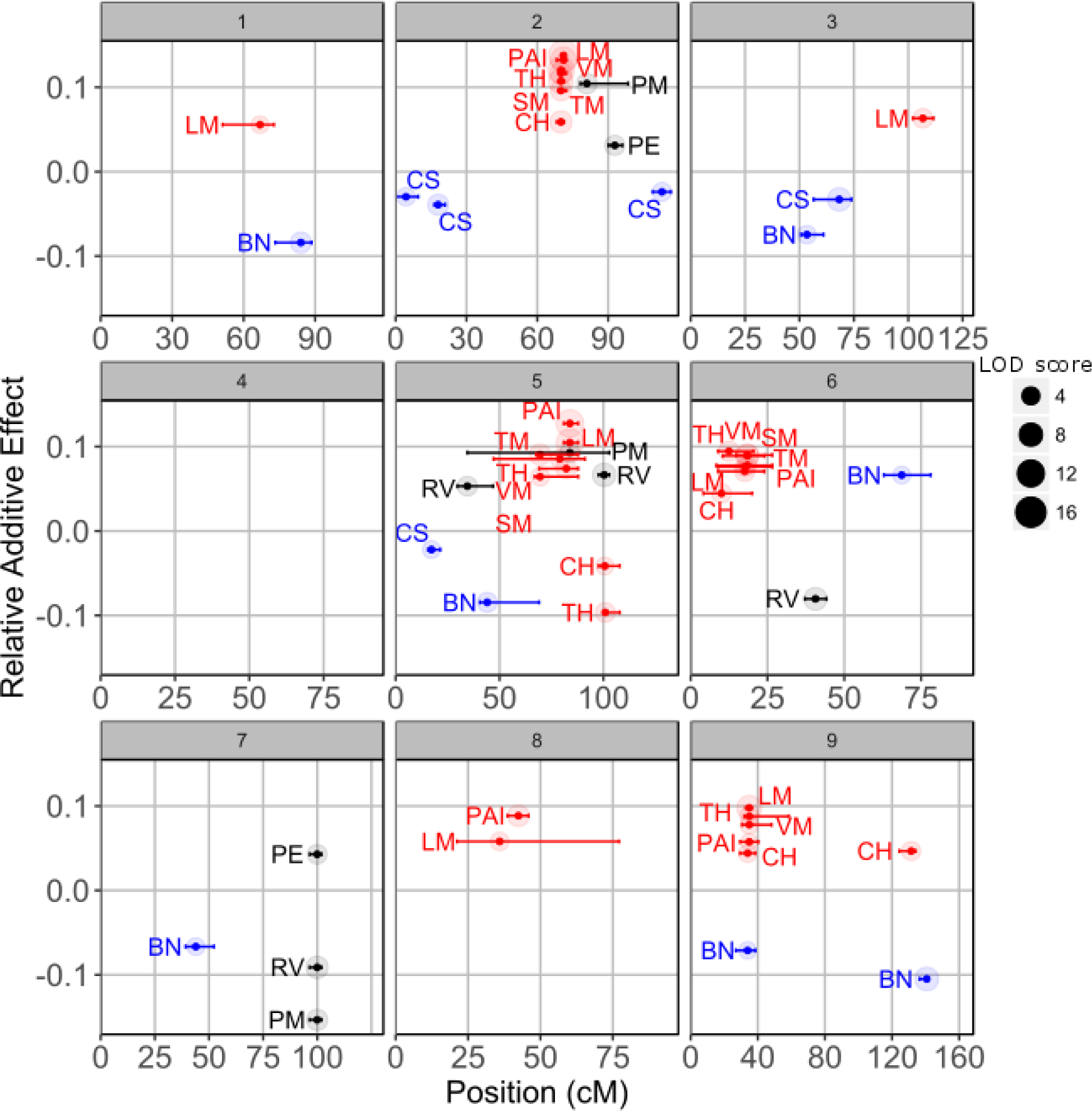
QTL mapping for PAI and directly measured biomass traits. Panels 1-9 correspond to each chromosome in the *Setaria* genome. Additive effects relativized by phenotypic mean are plotted against centimorgan position for each QTL. Error bars represent the 1.5 LOD score confidence intervals for each QTL’s location. Colored points represent QTL corresponding to biomass production (red) bushiness (blue) and partitioning of biomass to vegetative versus reproductive structures (black). PAI=plant area index, CS=clump spread, BN=branch number, CH=culm height, TH=second tiller height, LM=leaf mass per m^2^ ground, SM=stem mass per m^2^ ground, PM=panicle mass per m^2^ ground, VM=vegetative mass per m2 ground, TM=total mass per m^2^ ground, RV=reproductive to vegetative mass ratio, PE=panicle emergence days after sowing.

QTL for PAI were co-located with all four of the hotspots of QTL for traits related to above-ground biomass production evaluated in destructive harvests. These four hotspots found on chromosomes 2, 5, 8, and 9 featured QTL for between five and seven traits, including PAI, total mass, vegetative mass, leaf mass, stem mass, culm height, and tiller height. All PAI and biomass productivity QTL within the four hotspots had positive additive effects. Together, these loci appear to represent the primary features of the genetic architecture of above-ground biomass production in *Setaria*. There was very little overlap between the location of these QTL hotspots for biomass production and QTL for traits associated with bushiness or vegetative-to-reproductive biomass ratio. *S*ingle QTL each for PAI and leaf mass co-localized on chromosome 8 and both had positive additive effects. Additionally, two isolated QTL for leaf mass and culm height did not overlap with QTL for PAI. In contrast to all the other QTLs for traits associated with biomass production, QTL for culm height and tiller height had negative additive effects and co-located with a single QTL for reproductive-to-vegetative mass ratio (5@100). There was also very close correspondence between QTL identified for PAI and other productivity traits in the study with an independent field experiment on the same RIL population in Oklahoma^29^. This provides strong evidence in support of using HTP of PAI to evaluate the genetic architecture of field-grown grass crops biomass productivity.

Twelve QTL for traits related to plant bushiness and nine QTL for traits related to partitioning of biomass to vegetative versus reproductive structures were detected. Notably, there was minimal overlap between QTL for PAI and those for traits associated with plant bushiness and biomass partitioning rather than biomass production. So, while hemispherical imaging is a powerful tool for assessing the genetic architecture of productivity traits in the field, other more complex imaging techniques or laboratory based methods will be needed to quickly phenotype plant architectural traits^30^.

QTL for PAI captured the main features of the genetic architecture for directly measured traits related to biomass productivity (PC1), but in a non-destructive and far less laborious manner and without the need for intermediate calibration models. This was achieved with a minimal number of genotypes (186), replicates (1), and subsampling (2), confirming the method’s power to detect the genetic components of variation in biomass productivity. In addition to its high detection fidelity, hemispherical imaging has the characteristics of an ideal, scalable HTP technique. Compared to the manual traits that it parallels, hemispherical imaging is very efficient (4 person-hours for image collection and 7 for image analysis compared to 148 person-hours for destructive biomass harvest and 15 person-hours for sample weighing). The equipment used to collect and analyze hemispherical images is inexpensive and commercially available. This simplicity, low-cost and universally applicable principle of operation mean that the method could be deployed on diverse crops at any location by a small team of personnel with limited training. The equipment is compact and lightweight, meaning that it can also be deployed on a rover to further accelerate image collection, improve the signal-to-noise ratio, and uncover the temporal dynamics of biomass productivity.

## METHODS SUMMARY

### Validation experimental design

*Setaria viridis* and *Setaria italica*, the parents of the F7 RIL population, and the phenotypically intermediate RIL #161 were grown on the South Farms at the University of Illinois Urbana Champaign in summer 2014. The field site is rain-fed, tile-drained, has a deep, organically rich, Flanagan/Drummer series type soil. RIL #161 was selected as a phenotypic intermediate between *S viridis* and *S italica* because of its placement in the 50^th^ percentile for both culm height and tiller number. The experiment was a randomized complete block design with all three genotypes replicated in eight plots of which four were randomly chosen for measurement. Each plot was 4 m^2^ with 25 cm grid spacing between plants. Data was collected from an independent plot of each genotype on four dates through the growing season. This resulted in significant variation in height, biomass production, canopy architecture, and PAI. Measured plots were not used for subsequent data collection.

First, non-destructive estimates of PAI were generated using HemiView software (Delta-T Devices) to analyze 6 canopy hemispherical photographs taken either within or between planting rows near the plot center under diffuse light conditions (pre-dawn, dusk, or high cloud cover). Hemispherical photographs were taken with a GoPro Hero 3+ digital camera modified with a fully hemispherical lens and mounted on a miniature self-leveling gimbal. Second, 8 or 16 plants (depending on collection date) were harvested from each plot, and separated into leaf, stem, and reproductive tissues. Fresh leaves were laid flat and photographed with a digital SLR camera (Cannon EOS 7D, 50mm lens) alongside a scaling object to allow estimation of total leaf area using ImageJ (NIH). All tissues were then dried at 65°C and weighed.

### QTL experimental design

186 F7 recombinant inbred lines from an interspecific cross between *Setaria italica* x *Setaria viridis* were evaluated on the South Farms at the University of Illinois Urbana-Champaign in summer 2014. Seeds were germinated in greenhouses and transplanted by hand into a mechanically tiled field 7 days after sowing. The experiment was an un-replicated randomized design with six check plots for each parent. Data was collected from a single plot (36 plants, 1 m^2^, 20 cm grid spacing between plants) of each RIL and six plots of each parent genotype.

### Climate conditions

Over the duration of the validation experiment, the average air temperature was 21.74°C, the average humidity was 72.34%, and the cumulative rainfall was 37.23 cm. Over the duration of QTL experiment, the average air temperature was 19.37°C, the average humidity was 78.59%, and the cumulative rainfall was 18.85 cm.

### Phenotyping

Panicle emergence was measured as the number of days after sowing at which the panicle head was seen past the collar of the flag leaf. The angle between the outermost tillers (i.e. clump spread) was measured in the field with a modified protractor 50 days after sowing. Nondestructive estimates of PAI were generated using Hemiview software (Delta-T Devices) to analyze 2 canopy hemispherical photographs taken with a GoPro Hero 3+ digital camera either within or between planting rows at the plot center under diffuse light conditions at dusk or dawn when the canopy was near maximum size prior to senescence at the end of the growing season, between 67 and 70 days after sowing. The GoPro camera was customized by replacing the standard lens with a 1.39mm 190° fisheye lens and mounted on a self-leveling gimbal in order to insure the camera faced upwards from horizontal. The camera was consistently staged such that the top of the image was oriented north. Two identical camera setups were used and images were analyzed by two people. A common set of images were processed and analyzed to confirm a lack of camera or person bias. End of season destructive harvest was done on three representative center plants in each plot beginning 72 days after sowing. Plants were cut at the base, separated into leaf, stem, and reproductive tissues, and the following morphological traits were measured. Culm height was measured as the length from the base of the plant to the collar of the flag leaf on the first emerged tiller. Tiller height was measured as the length from the base of the plant to the collar of the flag leaf on the second emerged tiller. Basal circumference was measured with a length of twine wrapped around the root crown. Tiller number was measured as the count of tillers emerging from the bottommost node. Branch number was measured as the count of primary branches emerging from nodes one or higher. The separated leaf, stem, and reproductive tissues were dried at 65°C and weighed. Vegetative mass was calculated as the sum of leaf and stem mass. Total mass was calculated as the sum of leaf, stem, and panicle mass. All masses were standardized by planting density and are reported on a per unit ground area basis. The following R packages were used for data analysis and visualization: *ggplot2, plyr, reshape2,* and *ggrepel*. Data and scripts used in analyses is available in a .zip folder included in the supplemental information.

### Data transformation

Data were normalized using a second power, square root, or cube root transformation. Normality was assessed through the R function *shapiro.test* and the associated histograms and Shapiro Wilk’s values. Results of the transformation procedures are shown in Table S1.

### Trait correlations

Trait correlations were tested using the R function *cor* using pairwise deletion to generate Pearson’s coefficients of correlation and visualized with *corrplot*.

### Principle Component Analysis

Principle component analysis was performed using the R function *prcomp*, with default parameters. Only genotypes with complete sets of observations were used in the principle component analysis. Evaluation of individual eigenvectors for each trait was used to describe the significant PCs and used to parse the traits into biologically relevant groupings (Fig. 2). The three most significant principle components were treated as individual traits in correlation analysis.

### QTL analysis

QTL analysis was performed using “*foxy_qtl_pipeline”,* available at <u>https://github.com/maxjfeldman/foxy_qtl_pipeline,</u> written by Max Feldman and adapted from *R/qtl^31,32^*. QTL detection was done using forward-backward Haley-Knott regression in order to build a multiple QTL model from each trait. A genome scan interval of 1 cM and a window size of 10 were used. 1,000 permutations were performed to estimate LOD threshold values. Additive effects were estimated as half the distance between phenotypic averages for the two homozygotes. To compare additive effects across traits with different scales, additive effects were normalized as a percent of the phenotypic mean^33^. Co-localized QTL were grouped into “clusters” based on their mapping to same or neighboring markers where confidence intervals overlapped. Confidence intervals were calculated as the interval where the LOD score was within 1.5 units of its maximum. Lander and Botstein (1989) first proposed the use of 2 LOD support intervals and more recently Dupuis and Siegmund (1999) provided support for using the 1.5 LOD interval method^34,35^. The use of LOD support intervals as a method to estimate the location of QTL and define co-localized clusters continues in current plant biology QTL experiments^36^.

## Acknowledgements

Funded through Subaward No. 23009-UI, CFDA #81.049 between University of Illinois and Donald Danforth Plant Science Center Under Prime Agreement No. DE-SC0008769 from Department of Energy

We thank Scott Baker and Jarad Bear for constructing the camera gimbal. We thank Kannan Puthaval and Mac Singer for technical support at the SoyFACE research facility. We thank Marshall Alston- Yeagle, Kara Barto, Johnathon Yockey, Sarah Keeley, Zach Reynado, Finey Ruan, and the many project partners from the Danforth Plant Science Center, Carnegie Institute, Washington State University, and University of Minnesota that helped with transplanting seedlings.

---

**Table S1.** Summary of phenotyping results. Mean, number of observations, minima, maxima, range, fold change, and summary of transformation procedures for 14 measured traits are reported.

| Trait | Mean | N | Minimum | Maximum | Range | Fold change | Transformation | Wilks statistic | P-value |
| --- | --- | --- | --- | --- | --- | --- | --- | --- | --- |
| Basal circumference (mm) | 2607.51 | 195 | 40.11 | 8100.00 | 8059.89 | 200.94 | square | 0.98 | 0.0029 |
| Branch number (count) | 3.47 | 185 | 1.00 | 7.39 | 6.39 | 6.39 | square root | 0.99 | 0.1908 |
| Clump spread (degrees) | 3.93 | 203 | 2.99 | 4.89 | 1.90 | 0.64 | cube root | 0.99 | 0.6265 |
| Culm height (mm) | 20.20 | 188 | 10.25 | 27.47 | 17.22 | 1.68 | square root | 0.99 | 0.3258 |
| Leaf mass (g) | 9.07 | 189 | 4.41 | 16.69 | 12.28 | 2.79 | square root | 0.99 | 0.0459 |
| PAI (m<sup>2</sup>/m<sup>2</sup>) | 0.91 | 186 | 0.25 | 1.62 | 1.37 | 5.46 | square root | 0.98 | 0.0172 |
| Panicle emergence (days after sowing) | 39.17 | 210 | 29.00 | 49.00 | 20.00 | 0.69 |  | 0.93 | 0.0000 |
| Panicle mass (g) | 20.12 | 190 | 2.88 | 35.19 | 32.31 | 11.23 | square root | 0.99 | 0.4337 |
| Reproductive to vegetative mass ratio | 1.20 | 187 | 0.33 | 1.52 | 1.19 | 3.59 | square root | 0.93 | 0.0000 |
| Stem mass (g) | 14.22 | 191 | 6.87 | 22.49 | 15.62 | 2.27 | square root | 0.99 | 0.1684 |
| Tiller height (mm) | 359.68 | 187 | 113.67 | 721.67 | 608.00 | 5.35 |  | 0.99 | 0.1354 |
| Tiller number (count) | 1.81 | 187 | 1.26 | 2.69 | 1.43 | 1.14 | cube root | 0.99 | 0.0802 |
| Total mass (g) | 26.46 | 187 | 9.14 | 43.13 | 33.99 | 3.72 | square root | 0.99 | 0.5196 |
| Vegetative mass (g) | 16.92 | 188 | 8.44 | 28.01 | 19.57 | 2.32 | square root | 0.99 | 0.2685 |

**Table S2.**
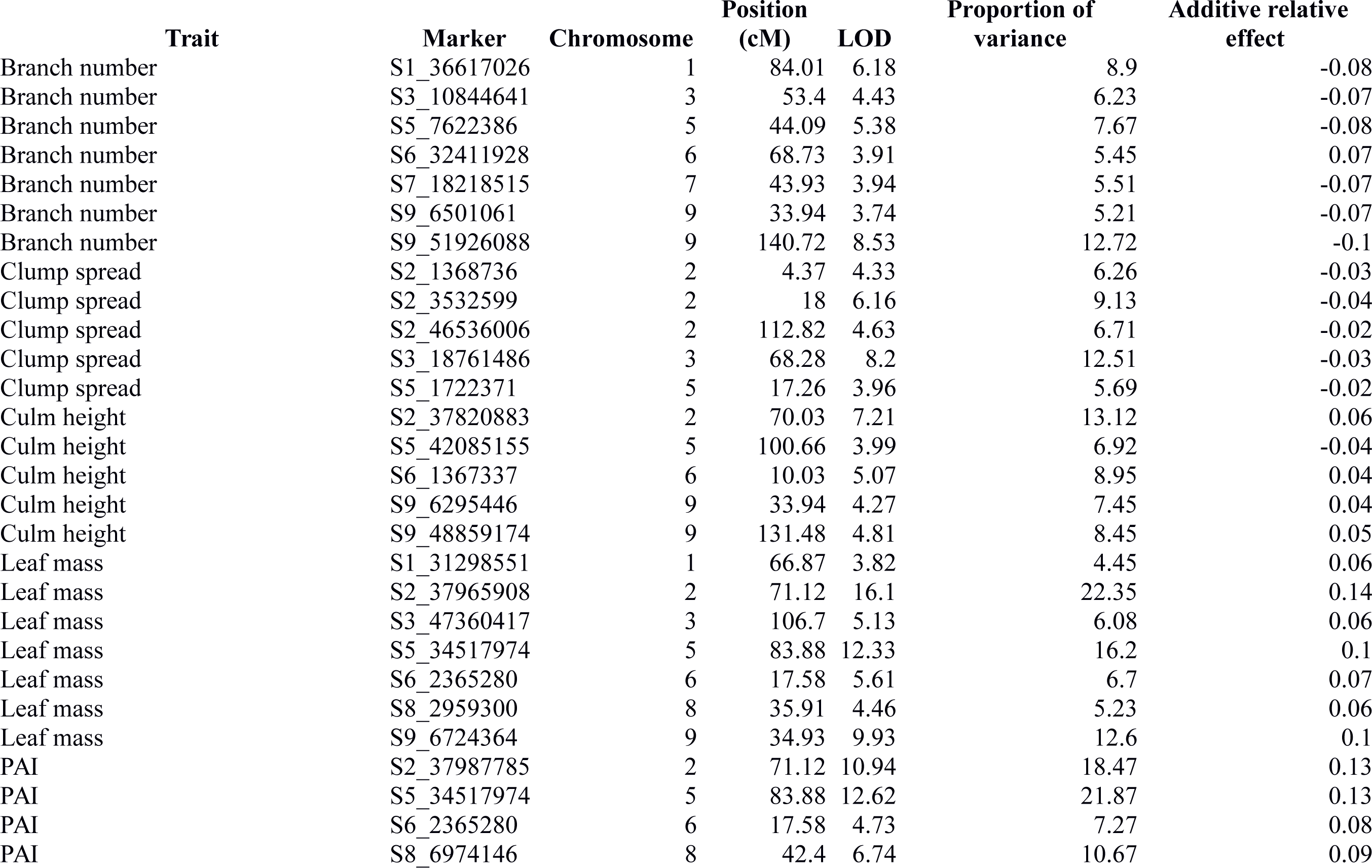
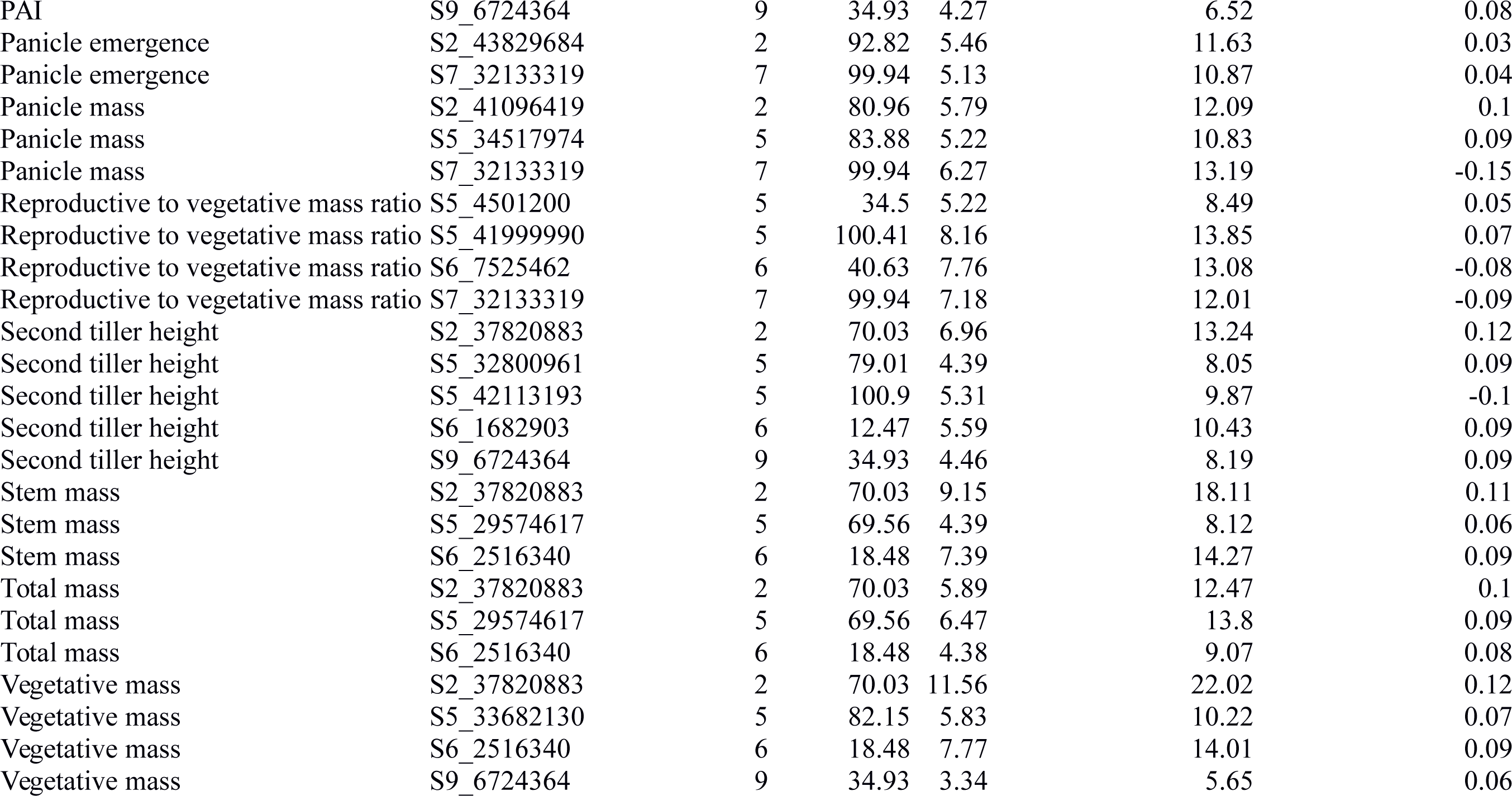
Summary of QTL results. Marker name, location, proportion of genotypic variance explained, and relative additive effect for QTL identified for 13 measured traits at a 0.05 detection threshold are reported.

| Trait | Marker | Chromosome | Position<br>(cM) | LOD | Proportion of<br>variance | Additive relative<br>effect |
| --- | --- | --- | --- | --- | --- | --- |
| Branch number | S1_36617026 | 1 | 84.01 | 6.18 | 8.9 | -0.08 |
| Branch number | S3_10844641 | 3 | 53.4 | 4.43 | 6.23 | -0.07 |
| Branch number | S5_7622386 | 5 | 44.09 | 5.38 | 7.67 | -0.08 |
| Branch number | S6_32411928 | 6 | 68.73 | 3.91 | 5.45 | 0.07 |
| Branch number | S7_18218515 | 7 | 43.93 | 3.94 | 5.51 | -0.07 |
| Branch number | S9_6501061 | 9 | 33.94 | 3.74 | 5.21 | -0.07 |
| Branch number | S9_51926088 | 9 | 140.72 | 8.53 | 12.72 | -0.1 |
| Clump spread | S2_1368736 | 2 | 4.37 | 4.33 | 6.26 | -0.03 |
| Clump spread | S2_3532599 | 2 | 18 | 6.16 | 9.13 | -0.04 |
| Clump spread | S2_46536006 | 2 | 112.82 | 4.63 | 6.71 | -0.02 |
| Clump spread | S3_18761486 | 3 | 68.28 | 8.2 | 12.51 | -0.03 |
| Clump spread | S5_1722371 | 5 | 17.26 | 3.96 | 5.69 | -0.02 |
| Culm height | S2_37820883 | 2 | 70.03 | 7.21 | 13.12 | 0.06 |
| Culm height | S5_42085155 | 5 | 100.66 | 3.99 | 6.92 | -0.04 |
| Culm height | S6_1367337 | 6 | 10.03 | 5.07 | 8.95 | 0.04 |
| Culm height | S9_6295446 | 9 | 33.94 | 4.27 | 7.45 | 0.04 |
| Culm height | S9_48859174 | 9 | 131.48 | 4.81 | 8.45 | 0.05 |
| Leaf mass | S1_31298551 | 1 | 66.87 | 3.82 | 4.45 | 0.06 |
| Leaf mass | S2_37965908 | 2 | 71.12 | 16.1 | 22.35 | 0.14 |
| Leaf mass | S3_47360417 | 3 | 106.7 | 5.13 | 6.08 | 0.06 |
| Leaf mass | S5_34517974 | 5 | 83.88 | 12.33 | 16.2 | 0.1 |
| Leaf mass | S6_2365280 | 6 | 17.58 | 5.61 | 6.7 | 0.07 |
| Leaf mass | S8_2959300 | 8 | 35.91 | 4.46 | 5.23 | 0.06 |
| Leaf mass | S9_6724364 | 9 | 34.93 | 9.93 | 12.6 | 0.1 |
| PAI | S2_37987785 | 2 | 71.12 | 10.94 | 18.47 | 0.13 |
| PAI | S5_34517974 | 5 | 83.88 | 12.62 | 21.87 | 0.13 |
| PAI | S6_2365280 | 6 | 17.58 | 4.73 | 7.27 | 0.08 |
| PAI | S8_6974146 | 8 | 42.4 | 6.74 | 10.67 | 0.09 |
| PAI | S9_6724364 | 9 | 34.93 | 4.27 | 6.52 | 0.08 |
| Panicle emergence | S2_43829684 | 2 | 92.82 | 5.46 | 11.63 | 0.03 |
| Panicle emergence | S7_32133319 | 7 | 99.94 | 5.13 | 10.87 | 0.04 |
| Panicle mass | S2_41096419 | 2 | 80.96 | 5.79 | 12.09 | 0.1 |
| Panicle mass | S5_34517974 | 5 | 83.88 | 5.22 | 10.83 | 0.09 |
| Panicle mass | S7_32133319 | 7 | 99.94 | 6.27 | 13.19 | -0.15 |
| Reproductive to vegetative mass ratio | S5_4501200 | 5 | 34.5 | 5.22 | 8.49 | 0.05 |
| Reproductive to vegetative mass ratio | S5_41999990 | 5 | 100.41 | 8.16 | 13.85 | 0.07 |
| Reproductive to vegetative mass ratio | S6_7525462 | 6 | 40.63 | 7.76 | 13.08 | -0.08 |
| Reproductive to vegetative mass ratio | S7_32133319 | 7 | 99.94 | 7.18 | 12.01 | -0.09 |
| Second tiller height | S2_37820883 | 2 | 70.03 | 6.96 | 13.24 | 0.12 |
| Second tiller height | S5_32800961 | 5 | 79.01 | 4.39 | 8.05 | 0.09 |
| Second tiller height | S5_42113193 | 5 | 100.9 | 5.31 | 9.87 | -0.1 |
| Second tiller height | S6_1682903 | 6 | 12.47 | 5.59 | 10.43 | 0.09 |
| Second tiller height | S9_6724364 | 9 | 34.93 | 4.46 | 8.19 | 0.09 |
| Stem mass | S2_37820883 | 2 | 70.03 | 9.15 | 18.11 | 0.11 |
| Stem mass | S5_29574617 | 5 | 69.56 | 4.39 | 8.12 | 0.06 |
| Stem mass | S6_2516340 | 6 | 18.48 | 7.39 | 14.27 | 0.09 |
| Total mass | S2_37820883 | 2 | 70.03 | 5.89 | 12.47 | 0.1 |
| Total mass | S5_29574617 | 5 | 69.56 | 6.47 | 13.8 | 0.09 |
| Total mass | S6_2516340 | 6 | 18.48 | 4.38 | 9.07 | 0.08 |
| Vegetative mass | S2_37820883 | 2 | 70.03 | 11.56 | 22.02 | 0.12 |
| Vegetative mass | S5_33682130 | 5 | 82.15 | 5.83 | 10.22 | 0.07 |
| Vegetative mass | S6_2516340 | 6 | 18.48 | 7.77 | 14.01 | 0.09 |
| Vegetative mass | S9_6724364 | 9 | 34.93 | 3.34 | 5.65 | 0.06 |

